# *BrrRCO*, encoding a homeobox protein, is involved in leaf lobe development in *Brassica rapa*

**DOI:** 10.1101/2023.08.02.551127

**Authors:** Pan Li, Tongbing Su, Yudi Wu, Hui Li, Limin Wang, Fenglan Zhang, Shuancang Yu, Zheng Wang

## Abstract

Leaf shape is a vital economic and developmental trait in leafy vegetable Chinese cabbage (*Brassica rapa* L. *subsp. pekinensis*), which varies from smooth to deeply lobed, whereas the molecular basis remains unclear. Here, we detected an incompletely dominant major quantitative trait locus (QTL) *qBrrLLA10* for the lobed-leaf trait in *B. rapa*, and identified *BrrRCO*, encoding a HD-Zip transcription factor, as the casual gene underlying lobed leaf formation in *B. rapa.* Genotyping analysis showed that abundant variations in promoter of *BrrRCO* is responsible for leaf lobe variation, and the expression levels and promoter activity were significantly affected by the promoter variations between two parents. BrrRCO was a nucleus-specific protein and possess highest expression level at the bases of each lobe. Ectopic overexpression of *BrrRCO* in *Arabidopsis* led to deeply lobed leaves have never been seen in wild type, and leaf lobe development was blocked when *BrrRCO* expression was down-regulated through virus-induced gene silencing assays. Taken together, our findings revealed that *BrrRCO* positively regulate leaf lobe formation in Chinese cabbage, and the cis-regulatory element modifications result in functional variation of *BrrRC*O, providing a novel insight into the leaf shape improvement in Chinese cabbage and other *Brassica* species.

**Highlight:** *BrrRCO* is essential for leaf lobe formation by repressing local growth of leaf margin cells in *B. rapa*, and cis-regulatory modifications cause the different allele effects.

## Introduction

Plant leaves, as the main source of photosynthetic product, play an important role in nutrition accumulation, gas exchange, nutrient distribution, water transport, and environmental adaption (Jiang *et al*., 2015). In different species, the function of leaves is relatively conservative, but leaf shape varies greatly, which indicate that leaf shape has been subject to natural selection during domestication and improvement (Wang *et al*., 2015). The variations of leaf shape mainly caused by the morphological changes of leaf margin, which varies from entire, serrated to deeply lobed (Nicotra *et al*., 2011). Deeply lobed-leaf is a type of leaf shape variation formed during plant adaptation to the natural environment. Compared with plant with entire leaves, plants with lobed leaves have higher photosynthetic efficiency and higher plant yield due to improved heat dissipation and canopy architecture by increased distance between leaves (Nicotra *et al*., 2011; Peppe *et al*., 2011).

Leaf development initiated at the flanks of shoot apical meristem (SAM), which is the source of all aboveground organs during the post-embryonic development of plants (Tsukaya *et al*., 2013). Initially, leaf primordia are initiated at the flanks of the shoot apical meristem. Subsequently, the basic morphology of the leaf, both leaf symmetry and the major subregions of the leaf including leaflet primordia, deeply lobe and serration, were established during the primary morphogenesis. Then, typical features of mature leaves, such as stomata and leaf hair, appeared through cell proliferation and differentiation during the secondary morphogenesis (Moon and Hake, 2011). Among them, the final leaf shapes are mainly determined by the meristem at the margin region of leaf primordium (termed as marginal blastozone), and localized enhancement and suppression of marginal blastozone growth eventually led to the marginal serrations, lobes, and leaflet formation (Dengler and Tsukaya, 2001).

Leaf lobe development and growth are regulated by many factors, such as hormone signaling, environmental signals, and complex genetic factors including transcription factor and microRNA (miRNA) and so on (Sluis and Hake, 2015; Sedivy *et al*., 2017; Satterlee and Scanlon 2019; Rowland *et al*., 2020). More recently, several genes regulating leaf lobe formation have been identified in model species, such as *Arabidopsis thaliana*, tomato (*Lycopersicon esculentum*), cotton (*Gossypium herbaceum* L.) and soybean (Glycine max) by using forward and reverse genetics, which involved in multiple gene families. The class I *KNOX* genes are mainly expressed in apical meristem to maintain meristem activity, and down-regulated in the initiated leaf primordia in *Arabidopsis*. The expression of *KNOXI* genes was repressed by *ASYMMETRIC LEAVES1/ROUGHSHEATH2/PHANTASTICA* (*ARP*) in *Arabidopsis*, resulting in simple leaves. In *Cardamine hirsuta*, *KNOXI* genes were reactivated and up-regulated in the leaves, resulting in the leaflet formation (Efroni *et al*., 2010). *NO APICAL MERISTEM/CUP-SHAPED COTYLEDON* (*NAM*/*CUC*) genes play an important role in regulating leaf margin development and leaflet formation. In *Arabidopsis,* three CUC members, *CUC1*, *CUC2*, and *CUC3*, were first discovered, and there is functional redundancy among these three genes. Overexpression of *CUC2* resulted in lobed cotyledon and serrated leaf margin that disappeared in *cuc2* mutant (Hasson *et al*., 2011). In tomato, the function loss of *NAM/CUC* genes leads to fewer and fused leaflets. Overexpression of *LAEFY* (*LFY*) in *C. hirsuta* resulted in more complex leaf morphology, with deepened leaf lobe and leaflets development at the petiole and axil (Monniaux *et al*., 2017). *LATE MERASTEM IDENTITY* 1 (*LMI1*), encoding a HD-Zip I transcription factor, was required in the formation of leaf serrations and bracts (Saddic *et al*., 2006). *REDUCED COMPLEXITY* (*RCO*) that arose from the gene duplication of its ancestral paralog *LMI1* was first discovered in *C. hirsuta,* and is required for leaflet formation (Vald *et al*., 2014). TCP genes repress leaf lobe development through reducing the auxin synthesis and polar transport and down regulating the *CUC* genes expression (Kant *et al*., 2009; Lucero *et al*., 2015; Koyama *et al*., 2017)

*Brassica* plants has a rich diversity of genetic and morphology, including leafy vegetables, Turnip, and oil crops. Leaf morphology varies enormously from smooth to deeply lobed in different *Brassica* crops, making it an ideal material for studying the mechanism leaf morphology evolution (Rast-Somssich *et al*., 2015). More recently, genetic basic and candidate genes of leaf lobe development were reported in some *Brassica* crops. In *B. napus*, the lobed-leaf trait was positively regulated by two tandemly duplicated *LMI1-like* genes underlying the incomplete dominant locus *BnLL1* (Ni *et al*., 2015; 2017; Hu *et al*., 2018; 2020). In *B. oleracea*, a semidominant gene *BoALG10* that encodes an alpha-1, 2-glucosyltransferase, was reported to regulate the feathered-leaved trait (Feng *et al*., 2020; 2022). Another two studies reported that *BoLMI1a,* the homolog of *LMI1*, has been proven to regulate the formation of lobed leaves in *B. oleracea* (Ren *et al*., 2019; Zhang *et al*., 2021).

*Brassica rapa* (*Brassica rapa pekinensis*. L) is an important leafy vegetable crops originated in China, and leaf shape is the target trait of natural and artificial selection during evolution (Kubo *et al*., 2010; Wang *et al*., 2015; Sedivy *et al*., 2017). Some natural populations with closely related genome vary considerably in leaf shapes, indicating more complex regulation mechanism of leaf shape development in *B. rapa*. Up to now, a quantitative trait locus (QTL) on the end of Chromosome A10 has been a research hotspot for leaf lobe formation (Nakao *et al*., 2010; Zhang *et al*., 2019), however, the underlying genes and their molecular regulation mechanism are largely unknown. Here, we identified *BrrqLLA10*, a major locus for lobe-leaf trait based on BSA-seq and traditional QTL mapping using DH and F_2_ populations, and revealed that *BrrRCO* was the causal gene underlying *BrrqLLA10* locus, which positively regulated the formation of leaf lobe, and sequence variations in *BrrRCO* promoter led to the leaf shape alteration in two parents, providing a novel insight to understanding the regulation mechanism of lobed-leaf development, thereby aiding genetic improvement of leaf shape trait in *B. rapa* and other *Brassica* species.

## Materials and methods

### Plant materials and mapping populations

MM is a European turnip line (*Brassica*. *rapa* L. ssp. *rapifera*) with deeply lobed leaves, which has been self-pollinated for six generations. BY is a Chinese cabbage inbred line (*Brassica. rapa* L. ssp. *pekinensis*) with serrated leaves. MM was crossed with BY to generate F_1_, which was used to produce a double haploid (DH80) population and F_2_ segregating populations (2018MB and 2021MB). The DH80, 2018MB and 2021 MB populations were planted in three filed trails in 2015, 2018 and 2021 at the central farm of Beijing Vegetable Research Center, Beijing, China, and used to conduct genetic analysis and QTL mapping.

To qualify the leaf lobe extent and complexity, we calculated the leaf dissection index of MM and BY as described previously (Li *et al*., 2023). The leaf lobe phenotype of individuals in DH and F_2_ populations was investigated on fifth leaf by visual inspection at seedling stage using a 1-3 scale:1= serrated leaf, 2=intermediate leaf, and 3=deeply lobed leaf, which were largely similar to BY, F_1_ and MM.

### Bulked segregant analysis by resequencing

To apply bulked segregant analysis (BSA) in our study, two extreme DNA bulks, the lobed-leaf bulk (LB-bulk, 50 lobed-leaf F_2_ individuals) and unlobed-leaf bulk (UL-bulk, 50 unlobed-leaf F_2_ individuals) were constructed using the 2018MB F_2_ population. The genomic DNA was isolated from the fresh leaves using modified cetyltrimethylammonium bromide (CATB) method (Murray and Thompson 1980). The DNA concentration was diluted to 100 ng/μL, and equal amount of DNA from 50 lobed-leaf and 50 unlobed-leaf F_2_ individuals were mixed to construct LB-bulk and UL-bulk. The sequencing libraries for two parent lines and two bulks were constructed and sequenced with an Illumina HiSeq 2500 (Oebiotech, China). Clean reads were obtained after filtered by the NGSQC toolkit (Dai *et al*., 2010) and aligned to the *B. rapa* reference genome V3.0 (Chen *et al*., 2022) using the software Burrows-Wheeler Aligner program (BWA) (Li and Durbin, 2009). Based on the alignment files, single nucleotide polymorphisms (SNPs) and insertion/deletion (InDels) were called using SAMtools (Sequence Alignment/Map Tools) (Li and Durbin 2009) and GATK (The Genome Analysis Toolkit) (McKenna *et al*., 2010). The software ANNOVAR was adopted to align and annotate the SNPs and InDels, for degerming the physical position of each SNP, (Wang et al., 2010). To identify candidate regions for lobed-leaf trait, the SNP-index for each pool and Δ (SNP-index) were calculated using the method previous reported (Li *et al*., 2020). The 95% confidence intervals were set as the screening threshold.

### QTL mapping for lobed leaf

Based on the genome resequencing data for two parents as our previous report (Zhang *et al*., 2018), molecular markers including SNP markers and small insertion/deletion (INDEL) were designed. Eventually, 586 SNP/InDel markers evenly distributed on the *B. rapa* genome were developed to genotype the individuals of the DH80 population. JoinMap 4.0 was employed to perform genetic linkage analysis based on the genotype data of DH80 population (Van Ooijen, 2006), and we calculate the map distances using the Kosambi mapping function (Kosambi 1943). The presence of QTLs was detected by MapQTL 5.0 software (Van Ooijen, 2004) via composite interval mapping (CIM) procedure with a threshold of LOD=3.0.

To further confirm and narrow down the QTL interval, we used two markers flanking the major QTL *qBrrLLA10* to genotype individuals of 2021MB population, and obtained 30 extreme recombinants. According to the comparison of resequencing data in candidate region between two parents, additional SNP markers were developed and used to genotype extreme recombinants, and narrow down the target interval of *qBrrLLA10*.

### Expression analysis of the candidate genes

Different sections of the fifth rosette leaf of MM and BY seedlings were sampled. Total RNA was extracted using TRIzol reagent (Invitrogen, Carlsbad, CA, USA). According to the manufacturer’s instruments, the SuperScript Reverse Transcriptase (Invitrogen) was used to synthesize the first-strand cDNAs. The expression patterns of candidate genes were explored by Real-time fluorescence quantitative PCR using a Roche thermocycler (LightCycler480, Roche, Switzerland). The *ACTIN2* (At3g18780) in *Arabidopsis* and *GAPDH* gene in Chinese cabbage were used as internal reference genes, and we calculated relative expression levels of candidate genes using the 2^−ΔΔCt^ method with three biological replicates (Livak *et al*. 2001). qRT-PCR primers were designed by Primer Primer 5, which are listed in Supplementary Table S1.

### Phylogenetic analysis of BrrRCO

The orthologs and paralogs of BrrRCO protein were obtained by blast against the *Brassica* database (BRAD, http://brassicadb.cn/), TAIR databases and National Center for Biotechnology Information (http://www.ncbi.nlm.nih.gov). We used MEGA 6.0 to perform multiple sequence alignments and phylogenetic tree construction through the neighbor-joining method with 1000 bootstrap replicates.

### Promoter activity analysis of *BrrRCO*

The promoter activity of *BrrRCO* in two parents were detected using the dual-luciferase (DUAL-LUC) reporter assay. The 1606-bp and 2735-bp segments of *BrrRCO* promoter from deeply lobed-leaf MM and unlobed-leaf BY were cloned into the transient expression vector CP461 to generate *BrrRCO^MM/BY^*-LUC under the control of *CaMV35S* promoter. These constructs were transformed into *A.tumefaciens* strain GV3101 and then injected into *Nicotiana benthamiana* leaves in different combinations with p19. After 48-96h, the luciferase (LUC) activity was analyzed by a Dual-Luciferase Reporter Assay System (Promega, USA).

### Subcellular localization of BrrRCO

The coding sequence (CDS) without termination codon was ligated into the binary vector *pSAT6-EYFP-N1* to construct *35S::BrrRCO-GFP* vector under the control of *CaMV35S* promoter. The recombinant *35S::BrrRCO-GFP* and positive control *35S::GFP* plasmids were transformed into the protoplast of Chinese cabbage. The green fluorescent protein (GFP) signals were subsequently observed using a Zeiss LSM 700 laser scanning confocal microscope (Germany).

### GUS staining

To explore the tissue-specific expression of *BrrRCO* in leaf development, we amplified the *BrrRCO* promoter sequences from MM and ligated into the *pMDC163* Gateway-compatible binary vector. The *proBrrRCO^MM^::GUS* plasmid was transformed into *Arabidopsis* wild-type plants. The seedlings of T_3_ transgenic positive lines were used as research material. The dissected samples of transgenic and controls plants were submerged in GUS staining buffer (Huayueyang, Beijing) at 37℃ overnight in darkness. Then treated tissues were then decolorized in 75% (v/v) ethanol for several times to remove the chlorophyll before observation.

### Functional analysis of *BrrRCO*

For overexpression assays, the full CDS of *BrrRCO* was amplified and inserted into the *KpnI*/*SpeI*-linearized binary vector *pCAMBIA1301* driven by *CaMV35S* promoter. The recombiant *35S::BrrRCO* and control vectors were separately transformed into wild-type *Arabidopsis* as described by Li *et al* (2023). The T_0_ seeds of transgenic plants were initially screened using 1/2 Murashige and Skoog (MS) medium containing 50 mg/L hygromycin. The positive transgenic plants were further confirmed through PCR amplification. The seedings of T_3_ generation positive lines were used for expression analysis and phenotype observation.

For knockout assays, a turnip yellow mosaic virus (TYMV)-induced gene silencing (VIGS) technique was used to knock-down *BrrRCO* as previous described (Li *et al*., 2023). A 40-nt exon sequence specific to *BrrRCO* was selected, and the corresponding 80-nt palindromic sequence with *SnaBI* restriction site was insert into the *SnaBI*-linearized pTY vector using a T4 DNA ligase. The recombinant pTY-BrrRCO and pTY-S control vector were introduced into *Escherichia coli* Stb13 for plasmid extraction in large quantity using OMEGA Plasmid Giga Kit D6920. Then the high concentration plasmids were diluted to suitable working concentration (300–500 µg/µl). For the first virus infiltration, about 2-4 fully expanded leaves of 2-3-week-old MM plants were used. The treated plants were cultured in darkness at 22℃ for 24h and then transferred to 22℃ under the 16h light/8h dark conditions. The second infection was carried out 7 days after the first, and 3-4 infusions were required to achieve silencing effect. The seedlings treated with empty pTY-S and ddH_2_O were used as a positive and negative control, respectively. The phenotype observation of newly grown leaves was conducted one week after the last infection, and the expression of *BrrRCO* gene in the aforementioned lines were detected.

## Results

### Phenotypic characterization and genetic analysis of lobed-leaf trait

Most turnip cultivars have lobed leaves, while Chinese cabbage cultivars have entire or serrated leaves. In this study, a European turnip line MM with deeply lobed leaves and a Chinese cabbage line BY with serrated leaves were chosen for phenotypic and anatomical comparisons (Fig. 1A, B). Examination with scanning electron microscopy (SEM) showed that cells of the leaf hydathodes (Fig. 1C) and leaf margin regions (Fig. 1D) are larger and typically long rod-like in unlobed-leaf BY. In lobed-leaf MM, cells of the leaf hydathodes (Fig. 1E) and base regions between two lobes (Fig. 1F) are smaller, round, and closely arranged.

**Fig. 1.**
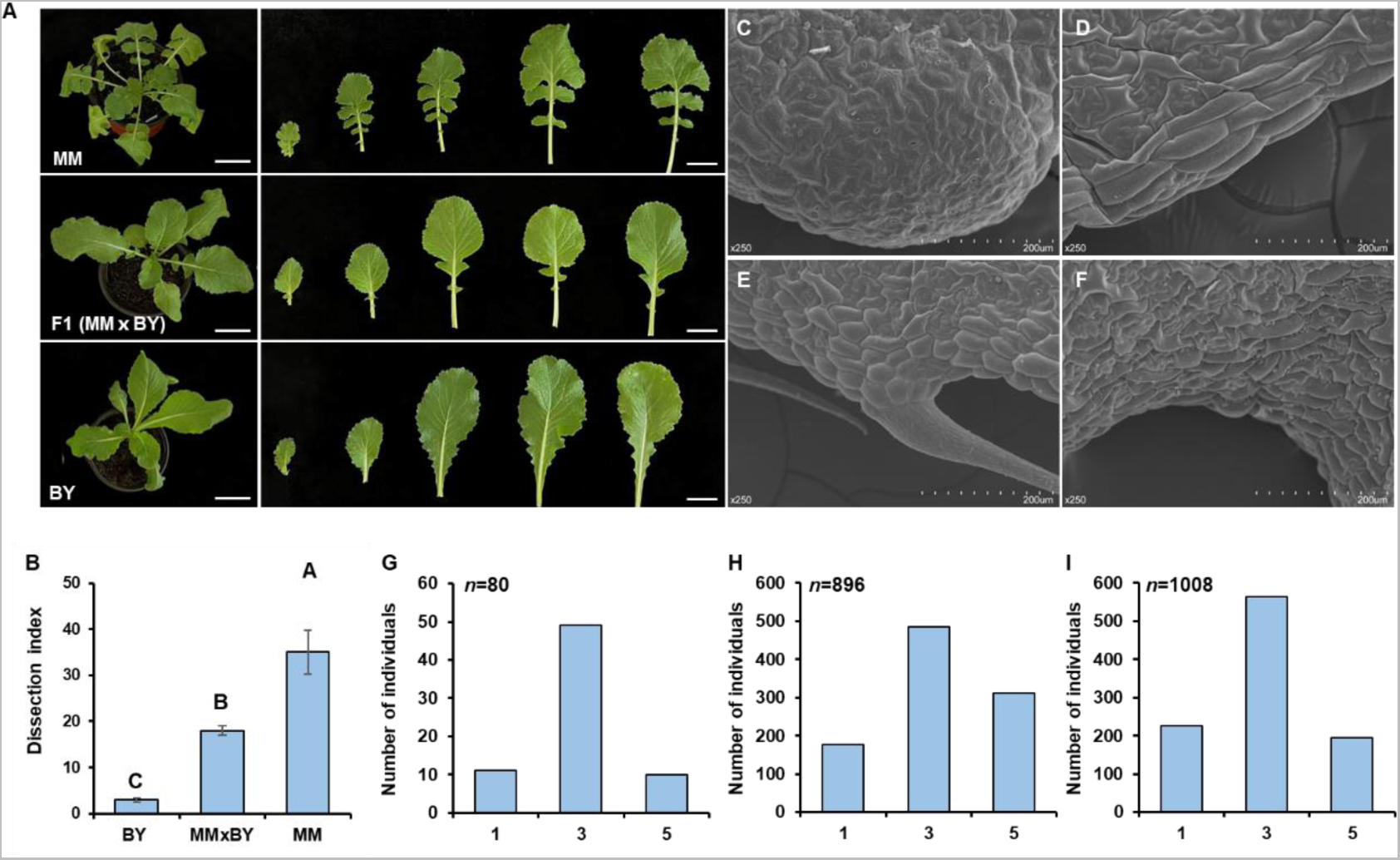
Phenotypes and physical characterization of leaf lobe in two parents and their F_1_ hybrids, and the frequency distribution of lobed-leaf trait in DH and F_2_ populations. (A) Representative leaf shape images of the two parents and their F_1_ hybrids. (B) Leaf dissection indices of two parents and F_1_ hybrids (n = 20). (C to F) Scanning electron micrographs of leaf margin cells in serrated-leaf BY (C, D) and lobe-leaf MM (E, F). (G to I) Frequency distribution of lobe-leaf trait in DH (G) and F_2_ populations. (H) MM × BY F_2_ population in spring of 2018 (2018MB). (I) MM × BY F_2_ population in spring of 2021 (2021 MB) MM: lobed leaf line. BY: entre leaf line. Bars represent the mean ± SD (n = 20). The scale bar is 1 cm.

To elaborate the genetic pattern underlying lobed-leaf trait, we crossed the deeply lobed-leaf MM and serrated-leaf BY. All the F_1_ plants showed semi-lobed leaves like those of the lobed-leaf parent MM, indicating the lobed-leaf trait is incompletely dominant (Fig.1A). Subsequently, one DH population (namely DH80) and two F_2_ segregating populations (namely 2018MB and 2021MB, respectively) were constructed. The leaf lobe frequency distribution in DH and two F_2_ populations were continuous and showed a largely normal distribution, implying the lobed-leaf trait is controlled by a quantitative trait locus from MM (Fig.1G, H, I).

### Identification of QTLs controlling leaf shape by BSA-seq and traditional QTL mapping

To identify the QTL controlling the lobed-leaf trait, BSA-seq was performed based on two extreme DNA bulks, lobed-leaf LB-bulk and unlobed-leaf UL-bulk. A total of 38 666 102 (13.42 average sequencing depth) and 36 148 178 (13.78 average sequencing depth) clean reads were generated from MM and BY, respectively. After filtering, 211 274 738 (63.72 average sequencing depth) and 194 522 146 (60.01 average sequencing depth) clean reads were obtained for LB and UL bulks, and mapped to the *B. rapa* reference genome ‘Chiifu-401-42’V3.0, respectively (Chen *et al*., 2022). About 1 282 540 SNPs and 576 671 INDEls were identified between the parental lines and the bulks. High-quality SNPs and INDELs were used to calculate the SNP-index and Δ (SNP-index) between the LB and UL bulks. The average SNP-index of each bulk was calculated in a 1-Mb interval using a 10-kb sliding window and was plotted against genome positions. Two statistically significant peaks in the plot of Δ (SNP-index), approximately 6.86Mb region (0-6.86Mb) on chromosome A02, and 1.15Mb (19.58-20.73Mb) on chromosome A10, were consider to be the candidate regions associated with leaf lobe formation and designated as *qBrrLLA02* and *qBrrLLA10* (*Brassica. rapa* L. ssp. *rapifera* lobed leaf A02 and A10) (Fig.2A).

**Fig. 2.**
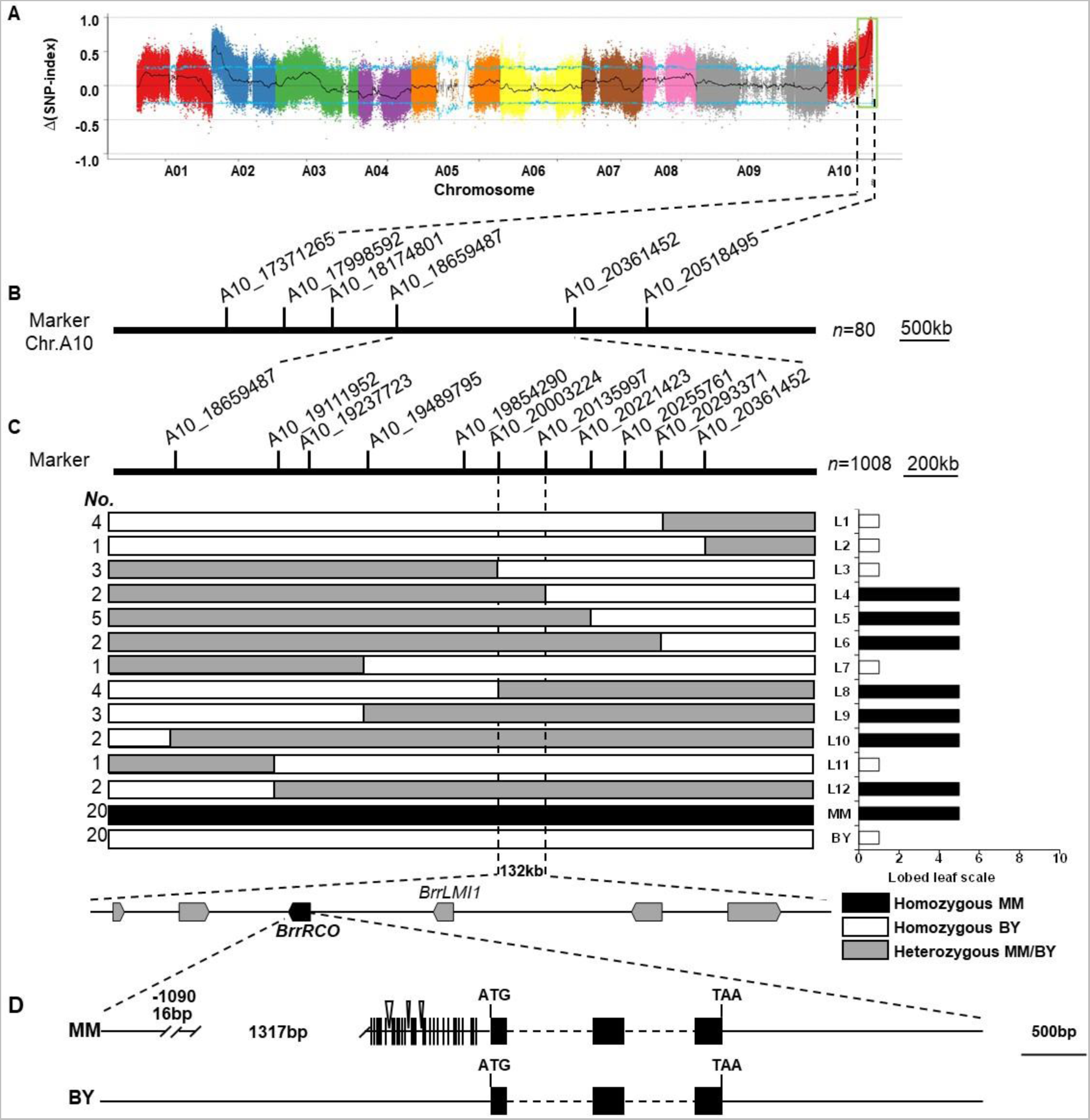
Map-based cloning of the *BrrRCO*. (A) Bulked-segregant analysis (BSA)-seq for the identification of leaf lobe related locus. (B) High-resolution linkage map of the *qBrrLLA10* region on chromosome A10. (C) The *qBrrLLA10* locus was delimited to a 132-kb region bounded by SNP markers A10_20003224 and A10_20135997. On the left, *No.*: number of recombinants in each type; (D) Gene structure of *BrrRCO* and sequence variation between MM and BY. The double slash and inverted triangle represent nucleotide deletion and insertion in the promoter region. The size of the exons and introns can be estimated using the scale bar at the bottom.

To confirm the results obtained by BSA-seq, classical QTL mapping with the DH80 population was also conducted. A major QTL *BrrqLLA10* (LOD=8.01) explaining 40.9% of the phenotypic variance was detected, which was located on the 18.65-20.36Mb region flanking by SNP markers A10_18659487 and A10_20361452 on the chromosome A10 (Fig. 2B; Supplementary Fig. S1, Supplementary Table S2). The physical location of *BrrqLLA10* was almost identical to that of BSA-seq analysis, supporting the existence of the major QTL locus *qBrrLLA10* of leaf lobe on chromosome A10.

To fine map the *qBrrLLA10*, the individuals of larger MM×BY F_2_ population (2021MB, 1008 individuals) were genotyped using two flanking SNP markers A10_18659487 and A10_20361452, and 30 extreme recombinants were obtained (Supplementary Table S3). Using nine newly developed SNP polymorphic markers, the extreme recombinants were genotyped and ultimately delimited the *qBrrLLA10* locus to a 132-kb interval bounded by SNP markers A10_20003224 and A10_20135997 on the chromosome A10 (Fig. 2C).

### Candidate genes mining for qBrrLLA10 locus

In the 132-kb interval of *qBrrLLA10*, 23 genes were predicted based on the *B. rapa* reference genome (Table S4). Among these genes, two tandemly duplicated genes (*BraA10g032440.3C* and *BraA10g032450.3C*) caught our attention based on their comparative gene annotation*. BraA10g032440.3C* (hereafter *BrrRCO*) was the homologue of the *RCO* in *Cardamine hirsuta*, which underlies in the leaflet formation (Vlad *et al*., 2014). *BraA10g032450.3C* (hereafter *BrrLMI1*) was the homologue of *LMI1* in *Arabidopsis,* playing a crucial role in the formation of leaf serration and bract (Saddic *et al*., 2006) (Supplementary Fig. S2).

We further investigated the expression levels of *BrrRCO* and *BrrLMI1* in different sections of fifth rosette leaves from MM and BY plants using qRT-PCR. The expression levels of *BrrRCO* in leaf base and leaf margin of MM leaves were both significantly higher than that in BY leaves, with the highest expression at the base of each lobe (Fig. 3A, B), whereas *BrrLMI1* expression was significantly lower in lobed-leaf MM than that of unlobed-leaf BY (Supplementary Fig. S3). Since *LMI1* and its homologs have been proposed to be a positive regulator of serrated leaf in *Arabidopsis* and other plants such as *GhLMI1-D1b* in upland cotton (Andres *et al*., 2017), *HcLMI1* in kenaf (Zhang *et al*., 2020) and *BrLMI1* in non-heading Chinese cabbage (Li *et al*., 2023), *BrrLMI1* should not be the candidate gene for *qBrrLLA10.* Therefore, *BrrRCO* was regarded as the most possible candidate gene for leaf lobe formation from MM.

**Fig. 3.**
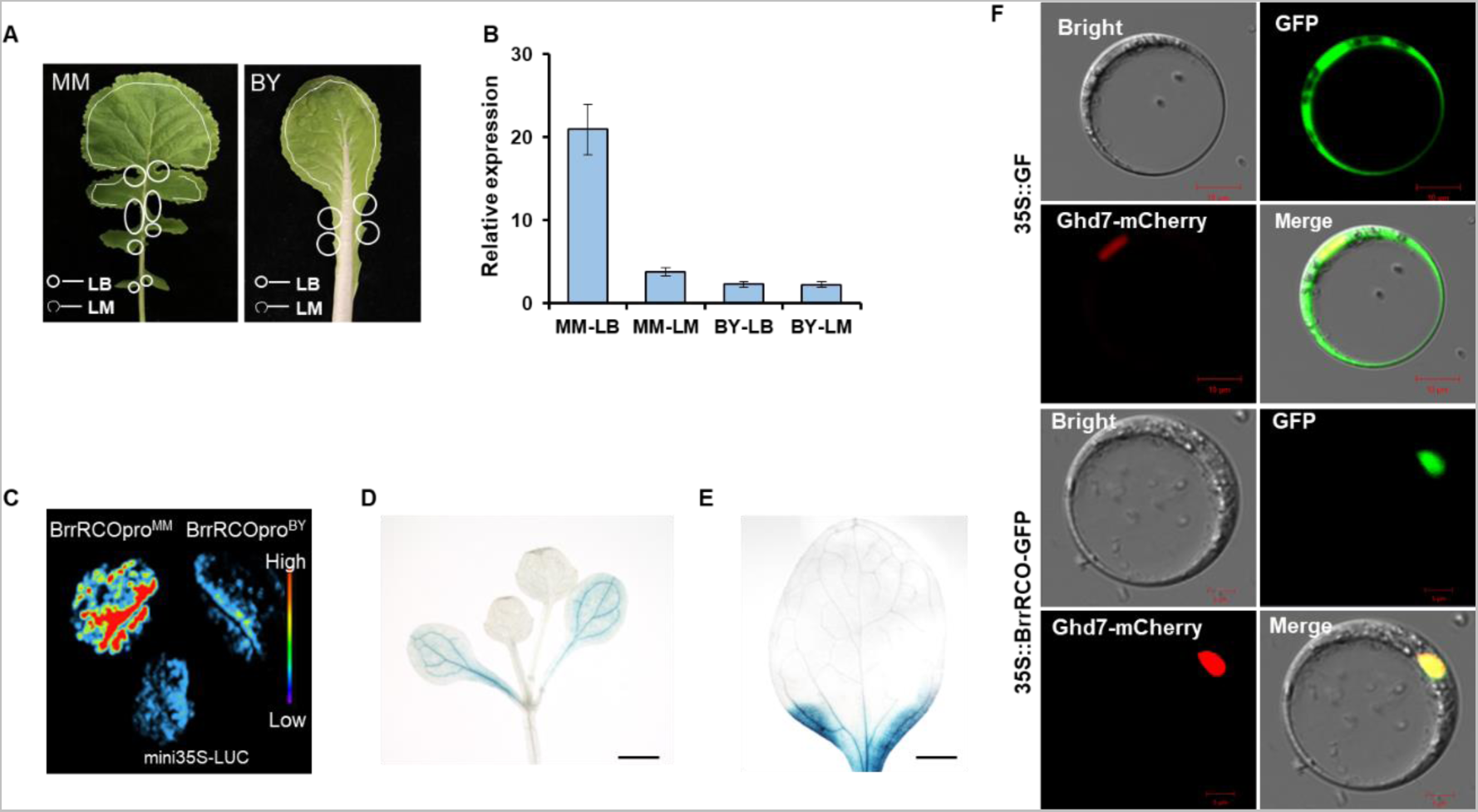
Expression patterns of *BrrRCO*. (A) Different leaf segments sampled for qRT-PCR analysis in MM and BY. (B) qRT-PCR analysis of *BrrRCO* expression in different leaf segments of MM and BY. (C) Transient dual luciferase (dual-LUC) assays of BrrRCO^MM^ and BrrRCO^BY^ in tobacco leaves. (D, E) Gus histochemical staining assays in *proBrrRCO::GUS* transgenic *Arabidopsis.* Representative histochemical staining in two-leaf stage seedlings (D, Scale bar: 500μm) and mature leaves (Scale bar: 0.1 cm) (F) Subcellular localization of BrrRCO in *B. rapa* protoplasts.

### Promoter polymorphisms in BrrRCO underlie its allelic variations

The nucleotide polymorphism of *BrrRCO* were further analyzed between two parental lines. The comparative sequence analysis of *BrrRCO* promoter sequence (∼ 3-kb), the entire gene and 3′flanking region (∼ 2-kb) showed that there is no difference in the coding region of BrrRCO between two parents, while a larger 1317-bpdeletion, a 16-bp deletion, 3 small InDels and 29 SNPs were detected within the *BrrRCO* promoter sequence of MM compared with BY and reference genome (Fig. 2D; Supplementary Fig. S4). Therefore, we speculated that promoter variations of *BrrRCO* is associated with alteration of expression level and leaf lobe formation. To test this, we tested the promoter activity of *BrrRCO* in MM and BY by using a dual-luciferase (DUAL-LUC) assay. A 1606-bp fragment (−1606 to −1) containing the 1317-bp deletion and 16-bp deletion from MM and a 2735-bp fragment (−2735 to −1) from BY were amplified and ligated to the linearized transient expression vector CP461, respectively. As shown in Fig. 3C, the *BrrRCO* promoter activity in deeply lobed parent MM was significantly higher than that in the serrated parent BY. We then randomly selected 12 different turnip lines with lobed leaves and 65 lines with unlobed leaves to investigate the promoter variation of *BrrRCO*. The result indicated that the 736-bp insert was present only in the lines with lobed leaves (Supplementary Fig. S5). These results indicated that the promoter variations of *BrrRCO* played a vital role in determining leaf morphology in *B. rapa*.

### BrrRCO tissue-specific expression and subcellular localization

To better understand *BrrRCO* tissue-specific expression patterns, we generated transgenic plants harboring a GUS reporter driven by *BrrRCO* promoter (*proBrrRCO::GUS*) to analyze tissue specific expression of *BrrRCO*. Histochemical analysis revealed *BrrRCO* expression is restricted to developing leaves, in the small regions at the base of leaf blade, and is absent from the meristem leaf boundary (Fig. 3D, E). According to previous reports, *LMI1* is expressed mainly in the leaf margins, serrations and in stipules and flowers, but do not accumulate at the leaf base (Saddic, *et al*., 2006; Daniela Vlad, *et al*., 2014; Li *et al*., 2023). Thus, we speculated that *BrrLMI1* and *BrrRCO* expressed in a near-complementary pattern to govern the leaf shape in *Brassica rapa*.

Subcellular localization of BrrRCO was performed by transiently expression of recombinant *p35S::BrrRCO* in the protoplasts of Chinese cabbage. The fluorescent signals of *p35S::BrrRCO* was exclusively localized in the nucleus (Fig. 3F).

### Verification of the function of BrrRCO in regulating lobe-leaf formation

As expression analysis indicated the lobed-leaf phenotype may be caused by the elevated expression of *BrrRCO*, we overexpressed the coding sequence of *BrrRCO* driven by *CaMV 35S* promoter into wild-type *Arabidopsis*, and 8 independent T_2_ transgenic lines were obtained. qRT-PCR assays indicated the expression levels of *BrrRCO* in three representative lines were increased significantly. *BrrRCO* overexpressing lines produced severely lobed-leaf never seen in the wild type, and the sinus region of rosette leaf was deepened (Fig. 4A-D), indicating that *BrrRCO* plays an important role in increasing leaf complexity in *B. rapa*.

**Fig. 4.**
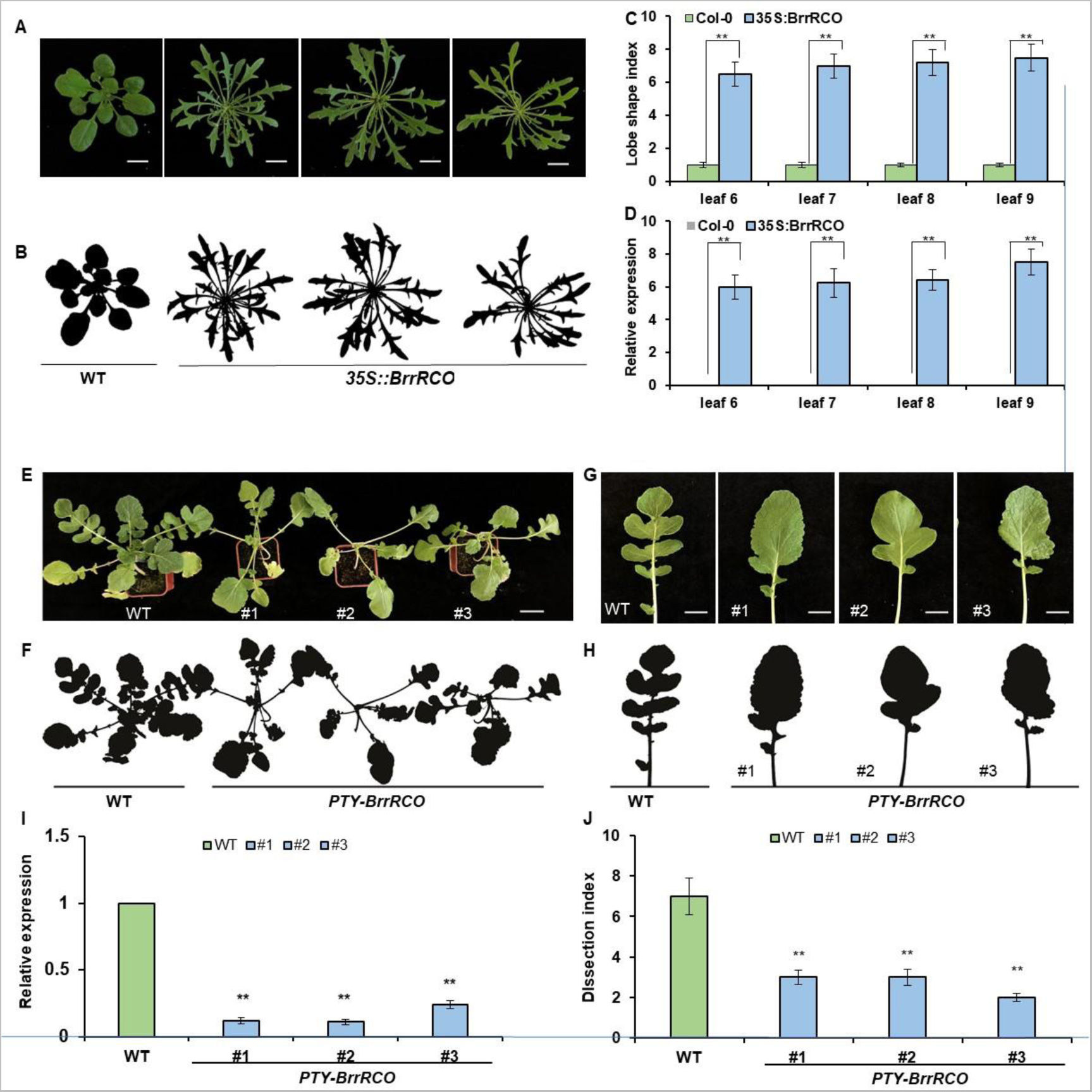
Functional analysis of BrrRCO in leaf lobe formation. (A, B) Leaf lobe phenotypes and silhouette of three independent T_2_ *BrrRCO* overexpression lines and Col-0 wild type *Arabidopsis* plants. Scale bars: 1cm. (C) Quantification of the number of serrations in plants shown in (A). (D) *BrrRCO* expression levels in rosette leaves 6 to 9 of overexpression and Col-0 plants. Error bars represent standard errors derived from three replications. (E to H) Leaf lobe phenotypes and silhouette after silencing of *BrLMI1* in MM. Scale bars: 2 cm. (I) *BrrRCO* expression level of in wild type and *BrrRCO*-silenced MM. Error bars represent standard errors derived from three replications. (J) Quantification of the number of lobes in plants shown in (G, H). Double asterisk denotes statistically significant difference to wild type in Student’s t-test.

To further validate the functions of *BrrRCO* in leaf lobe development in *B. rapa*, we use a VIGS technology to obtain loss-of-function lines. We inserted *BrrRCO*-specific exon target sequences into pTY-S plasmid to constructed the recombinant pTY-*BrrRCO* vector, which were inoculated into 2-3 fully expanded true leaves of lobed-leaf MM seedlings by friction inoculation. The seedlings treated with pTY-S were used as controls. The expression level of *BrrRCO* was significantly down-regulated in pTY-*BrrRCO* positive lines. MM seedlings with down-regulated *BrrRCO* expression produced a visible reduced lobed-leaf phenotype in newly-developed leaves compared with controls (Fig. 4E-J). These results indicated that *BrrRCO* positively regulates the lobed-leaf formation in *B. rapa*.

## Discussion

Leaf shape is an important commercial trait in leafy vegetables, and an ideal model for studying morphological evolution and phenotypic novelty (Mentink and Tsiantis, 2015; Rast-Somssich *et al*., 2015). Extensive morphological variation, including leaf lobe, exist in the *Brassica* species (Piazza *et al*., 2010). So far, several genes regulating the formation of leaf lobe have been verified in some *Brassica* species, such as *B. napus* (Ni *et al*., 2015; Ni *et al*., 2017; Hu *et al*., 2018; 2020), *B. oleracea* (Feng *et al*., 2020; 2022). However, the functional gene and molecular regulatory mechanism underlying leaf morphological diversity is still poorly understood in *B. rapa*. To address these issues, we identified a major QTL for leaf lobe formation, *qBrrLLA10*, by integrating BSA-seq and traditional QTL mapping, and further confirmed that *BrrRCO* was the causal gene underlying the *qBrrLLA10* locus, which encodes an HD-ZIP I transcription factor. The significantly up-regulated expression levels and higher promoter activity of *BrrRCO* in lobed-leaf parent MM indicated that the promoter modifications are vital for functioning of *BrrRCO* by enhancing its transcription and the up-regulated expression was responsible for deeply lobed-leaf development, which further confirmed that cis-regulatory divergence of *LMI1* and its homologous gene and consequent spatiotemporal modification of their expression is an evolutionary hotspot in leaf lobe development (Vuolo *et al*., 2016; Streubel *et al*., 2018; Hu *et al*., 2018, 2020; Zhang *et al*., 2021; Bo *et al*., 2022). However, we have not disclosed how promoter variations affect *BrrRCO* expression in two parents. A further exploration of upstream regulatory inputs that delimits *BrrRCO* expression may help us to identify the polymorphisms that are responsible for the *BrrRCO* transcription.

The relationship between the patterns of cell division and expansion and leaf shape has been a hotspot in leaf morphogenesis research. In *C.hirsuta*, *ChRCO* was reported to increase leaf complexity by repressing growth at the boundaries between leaflets/lobes via regulating genes related with cytokinin (CK) (Vlad *et al*., 2014; Hajheidari *et al*., 2019). Recently, Wang *et al* (2022) reported a preferred developmental path created by synergy between *RCO* and *KNOX1* genes, in which, *KNOX1* regulated prolonged and anisotropic cell growth in the basal domain of leaves, combined with local growth repression by *RCO*, contributes to leaflet formation. In our study, compared with the larger and rod-shape cells of leaf marginal regions of serrated-leaf parent BY, the smaller and compact cells of the sinus and marginal regions of deeply lobe-leaf parent MM indicated that *BrrRCO* repressed the local cell growth of leaf margins in *B. rapa*. In addition, BrrRCO protein was localized in the nucleus, which was consistent with function of *BrrRCO* as a transcription factor and indicated that *BrrRCO* may be involved in leaf shape diversity by regulating various downstream target genes. Therefore, to make the regulatory network more complete, mining the downstream target genes through which *BrrRCO* acts to repress growth will be needed in the future.

*RCO* was discovered in *C. hirsuta* based on its simple lobed leaf mutant phenotype, and *RCO*-type genes are also required for leaf shape diversification in the crucifer family. *RCO* arose by gene duplication of its ancestral paralog *LMI1* (Sicard et al., 2014; Vlad *et al*., 2014). *BrLMI1* was identified to be a positive regulator of leaf shape complexity, and cis-regulatory divergence is directly associated with the formation of leaf lobe in our previous study (Li *et al*., 2023). Here, we identified that *BrrRCO* is one of the causal genes regulating leaf lobe formation in *B. rapa* based on its expression patterns and functional analysis. To understand how *BrrRCO* and *BrLMI1* synergistically affecting the leaf shape diversity in *B. rapa*, we assayed the expression patterns between the two genes using reporter gene assays in expanding leaves of *A. thaliana.* The *BrLMI1* expression was defined in the leaf margin, with highest expression in the serrations, similar to its *A. thaliana* paralog (Saddic *et al*., 2006 Li *et al*., 2023). *BrrRCO* was expressed only at the base and the sinus of leaf blade as founded in *C. hirsuta* (Vlad *et al*., 2014). Based on the above findings, we speculated that *BrrRCO* and *BrLMI1*worked in a near-complementary pattern around the leaf margin growth to govern the formation of leaf lobe in *B.rapa*. which still needs further verification.

Gene duplication is a major driver for increasing biological complexity, which allow novel functions to evolve, together with functional divergence of newly duplicated paralogs (Fisher, 1935; Haldane, 1932). More recently, *LMI1/RCO* locus was proved to be a major driver for leaf shape diversification. In the *Brassicaceae*, this locus comprises a tandem gene triplication that have evolved through two independent gene duplication events. The gene in position 1, *LMI1*, retaining most of the ancestral function, is a regulator of inflorescence meristem identity and leaf serration and bract formation in *A. thaliana* (Sicard *et al*., 2014; Vlad *et al*., 2014; Saddic *et al*., 2006). The gene in position 2, *RCO*, which is required for leaflets formation in the compound leaf species *C. hirsuta*, arose from the gene duplication of its ancestral paralog *LMI1* after the divergence of *Aethionema* and before the last common ancestor of *Arabidopsis* and *Brassica* (Vald *et al*., 2014). The gene in position 3, *LMI1-like 3*, arosing from *RCO* duplication, is not expressed in *C. hirsute* (Vlad *et al*., 2014). The secondary loss of RCO-type genes contributes to the evolution of simple leaves in *A. thaliana.* In *B. rapa* genome, four *RCO/LMI1*-type genes were detected based on a genome-wide search for homologues of BrrRCO. To better understand the gene ancestry of *RCO/LMI1* locus in *B. rapa*, the synteny relationship and sequence homology among RCO/*LMI1* paralogs from *B. rapa*, *A. thaliana*, *A. lyrata*, and *C. hirsute* were reanalyzed (Fig. S2). The maximum-likelihood phylogenetic tree splits into two well-supported clades. Consistent with previous report (Vlad *et al*., 2014), all the *LMI1* genes clustered in the first clade, and the *RCO* and *LMI1-like 3* gene were grouped in a second clade, which reflecting the gene duplication events that occurred at this locus. In addition, the absence of *LMI1*-*like3* genes indicated that *RCO* duplication did not occur after the split of *RCO* from *LMI1* in *B. rapa* genome.

In addition to *qBrrLLA10*, another QTL *qBrrLLA02,* located in a 6.86Mb region (0-6.86Mb) on chromosome A02 was also detected. Coincidentally, the physical location of a *LMI1-like* gene, *BraA02g001070.3C* (hereafter *BrrA02.LMI1*), obtained through a genome-wide search for *BrrRCO* homologues in *Brassica* database, was in the interval of *qBrrLLA02.* Therefore, we speculated that *BrrA02.LMI1* also affect leaf lobe formation in MM. Is this gene involved in regulating leaf lobe formation in *B. rapa*? What is the molecular mechanism of this gene on leaf shape diversity? To address this question, further fine mapping and functional identification of this gene will be needed in the future.

In summary, we revealed *BrrRCO* as a promising gene underlying leaf lobe formation in *B. rapa*, which worked in a near-complementary pattern with *BrLMI1* around the leaf margin growth to regulate leaf lobe formation. Cis-regulatory variation of *BrrRCO* altered its activity in the developing leaf and lead to the difference in leaf margin dissection.

## Acknowledgements

We are grateful to Dr. Jinghua Yang (Zhejiang University) for his help in VIGS assays.

## Author contributions

ZW, PL, and SY designed the experiments. PL, carried out the experiments. PL, and ZW performed data analysis. PL wrote the paper. HL and ZW revised the paper. TS, SY and ZW provided guidance to phenotype investigation assays. All authors discussed the results and commented on the manuscript.

## Data availability

The datasets supporting the conclusions of this article are included within the article and its supplementary data published online.

## Supplementary data

Fig.S1 QTL likelihood maps by multiple QTL model (MQM) on ten Chromosomes.

Fig.S2 Phylogenetic tree of BrrRCO and LMI1/RCO-like proteins from various species;

Fig.S3 qRT–PCR analysis of *BrrLMI1* expression in various segments of leaves in MM and BY.

Fig. S4 (A) Genomic sequence, (B) amino acid sequence and (C) promoter sequence alignment of *BrrRCO* in MM and BY genomes.

Fig. S5 The indel marker designed based on the deletion in the promoter of *BrrRCO* showing co-segregation with the leaf lobe trait in parents and different inbred lines.

Table S1 Primers used for fine-mapping, gene amplification, qRT-PCR, vector construction and the Indel markers

Table S2 Quantitative trait loci (QTL) for leaf lobe detected in DH populations

Table S3 Genotypes and phenotypes of the extreme recombinants used for fine-mapping of *BrrqLLA10*

Table S4 Annotated genes in the *BrrqLLA10* region between SNP markers A10_ 20003224 and A10_ 20135997

