## Supplemental figures for "*BrrRCO*, encoding a homeobox protein, is involved in leaf lobe development in *Brassica rapa*"

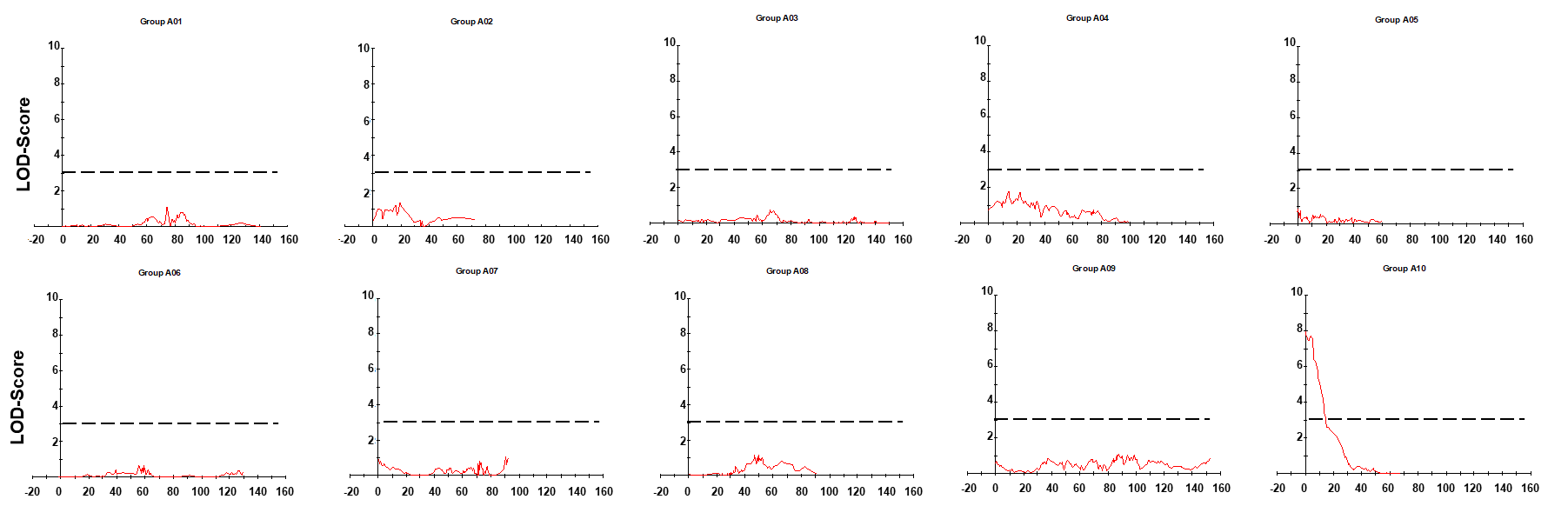

Fig.S1 QTL likelihood maps by multiple QTL model (MQM) on ten Chromosomes.

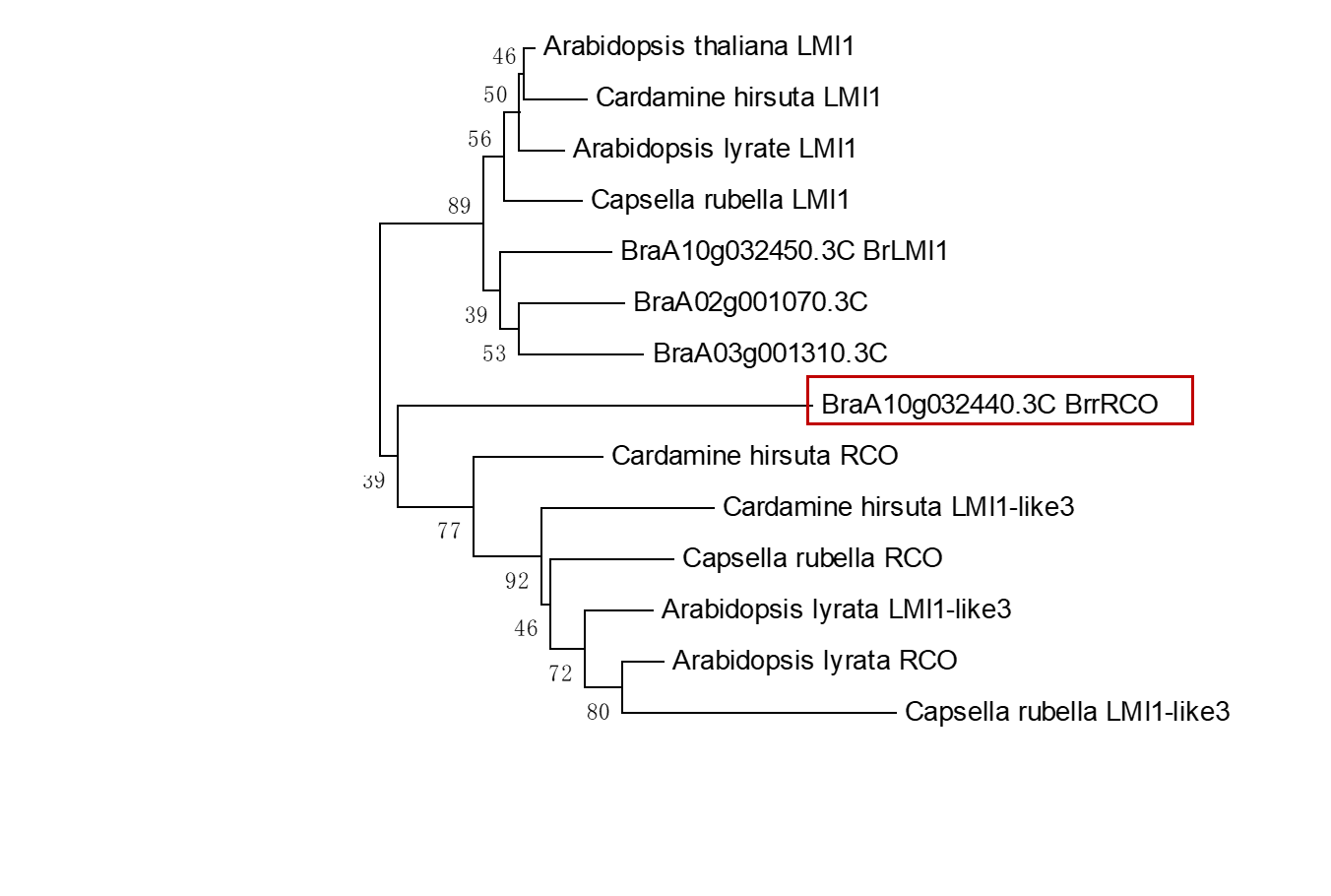

Fig.S2 Phylogenetic tree of BrrRCO and LMI1/RCO-like proteins from various species;

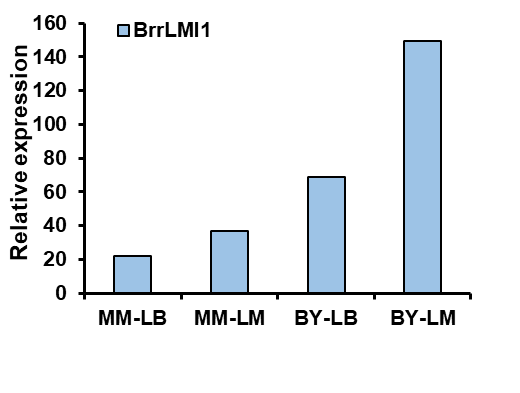

Fig.S3 qRT–PCR analysis of *BrrLMI1* expression in various segments of leaves in MM and BY.

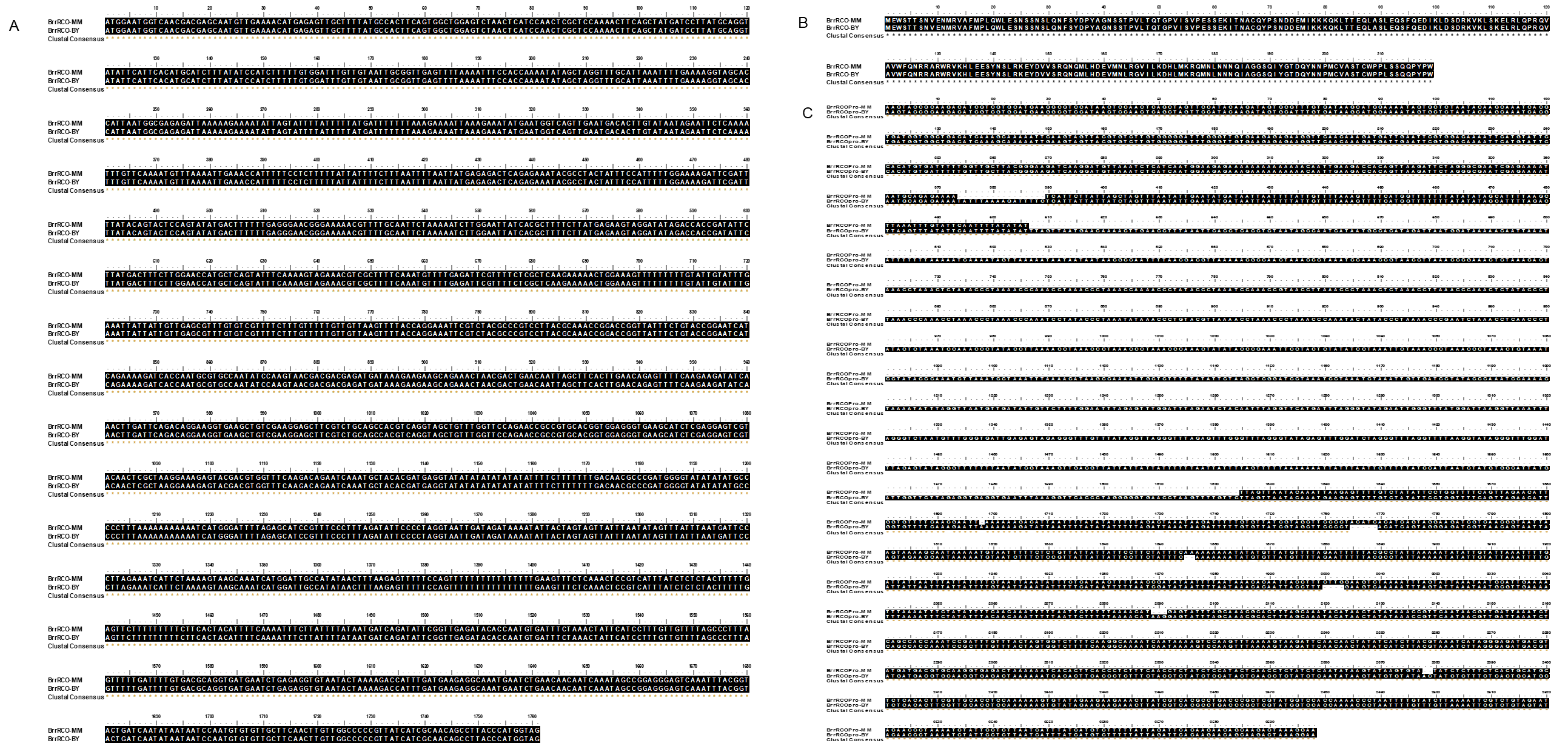

Fig. S4 (A) Genomic sequence, (B) amino acid sequence and (C) promoter sequence alignment of *BrrRCO* in MM and BY genomes.

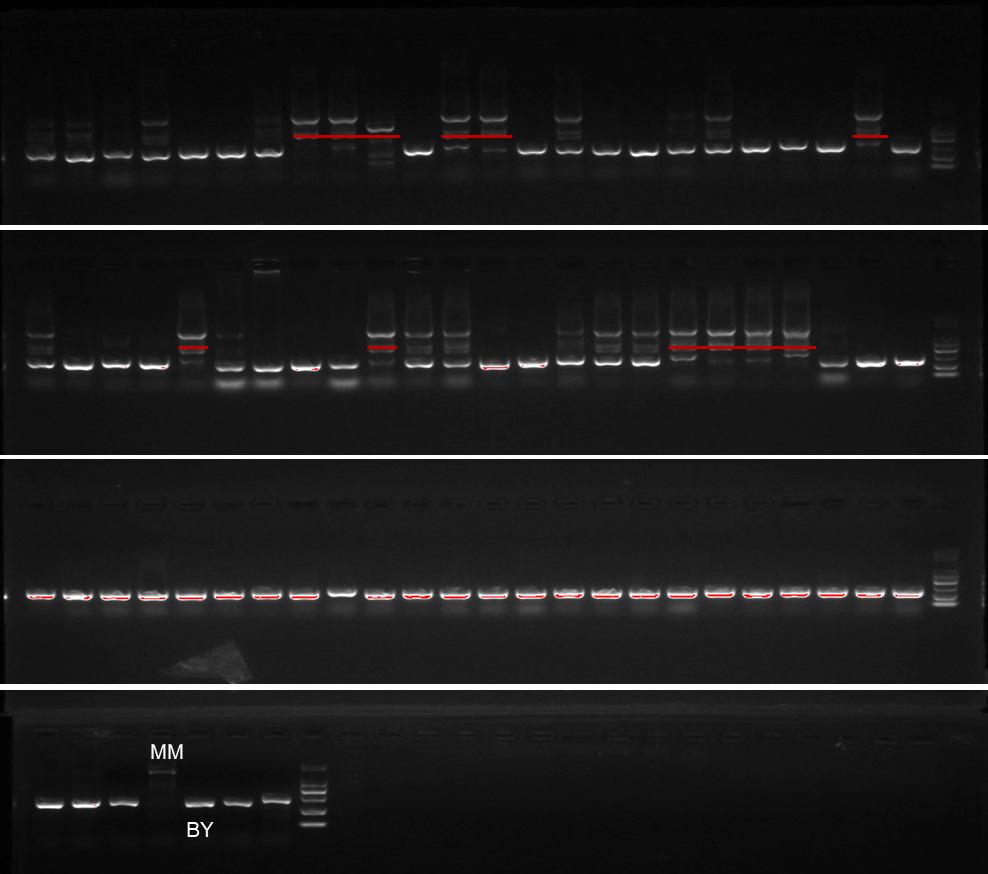

Fig. S5 The indel marker designed based on the deletion in the promoter of *BrrRCO* showing co-segregation with the leaf lobe trait in parents and different inbred lines.
